# A novel TRPA1 gain-of-function variant associated with painful sensory neuropathy acts through a PIP_2_ gating mechanism

**DOI:** 10.64898/2026.08.13.744390

**Authors:** Maddalena Comini, Tanadet Pipatpolkai, Sophia Clyde, Sanda Van Kruning Kodele, Matilde Laura, Andreas C Themistocleous, David LH Bennett

## Abstract

TRPA1 (transient receptor potential ankyrin 1) is a non-selective, calcium-permeable cation channel that mediates pain by detecting environmental irritants and thermal stimuli. Although the role of TRPA1 in modulating pain perception is relatively well established, so far only a few human TRPA1 variants (N855S and A172V) have been associated with inherited neuropathic pain disorders. Here, we describe a novel TRPA1 variant (p. M978V) identified in two human subjects presenting with painful sensory neuropathy. Electrophysiological recordings demonstrate that the M978V variant confers gain-of-function properties to the TRPA1 channel, especially in response to allyl isothiocyanate (AITC; mustard oil), a well-characterised TRPA1 agonist. The M978V substitution enhances current density and shifts the half-maximal activation potential, rendering the channel more readily activated by electrophilic agonists, such as AITC. Furthermore, the mutant channel exhibits increased plasma membrane expression following AITC stimulation, suggesting that this single amino acid substitution affects both channel gating and trafficking. Using all-atom molecular dynamics simulation (MD), we highlighted that the variant is adjacent to the PIP_2_ binding site on the TRPA1 channel. We further show that depletion of the membrane phospholipid phosphatidylinositol 4,5-bisphosphate (PIP_2_) increases current density in both WT and M978V channels. Importantly, the gain-of-function phenotype conferred by the M978V variant in response to AITC is dependent on the presence of PIP_2_. Collectively, our findings provide further evidence supporting the role of TRPA1 in human painful channelopathies and identify a previously unrecognised PIP_2_-dependent mechanism that regulates TRPA1 gain-of-function.

**Significance Statement:** In this study we characterised the mechanism by which a rare TRPA1 variant leads to painful sensory neuropathy and discovered a novel modulatory PIP₂-mediated regulation. Our *in vitro* data show that the variant confers gain-of-function properties to TRPA1 by enhancing its current density and open probability, as well as the channel’s surface membrane expression, in response to AITC, a known TRPA1 agonist. We also identified a novel interaction site for PIP_2_, a modulatory anionic lipid in the membrane of TRP channels. We have shown that abolishing endogenous PIP_2_ facilitates TRPA1 channel activation and that PIP_2_ is necessary for the variant’s gain-of-function properties, highlighting a new potential therapeutic avenue for neuropathic pain disorders.

## Introduction

TRPA1 is the only mammalian member of the TRPA family [1]. It is expressed in sensory neurons and has been implicated in nociception (through its response to environmental irritants), itch mechano- and thermo-sensation [2, 3]. Mice lacking TRPA1 display altered pain behaviour and show behavioural deficits in response to common TRPA1 agonists, such as mustard oil (AITC), menthol, cold temperatures [4, 5], and punctuate mechanical noxious stimuli [6]. TRPA1 also plays an important role as a sensor for cell death and cellular damage [7, 8], and is involved in ROS tolerance in cancer cells [9, 10]. At molecular level, activation of sensory TRP channels leads to cell depolarisation and subsequent increase of intracellular Ca^2+^ [11], as well as release of neuropeptides, like substance P and CGRP [12], which in turn promotes protective responses such as pain and inflammation. TRPA1 channels are largely expressed in primary nociceptors, where they respond to exogenous compounds (such as AITC and menthol, amongst others) as well as endogenous triggers (e.g., oxidative stress and inflammatory mediators), and have been linked to several human pain disorders, such as the familial episodic pain syndrome, firstly identified in a Colombian family [13]. In sensory neurons, when activated, TRPA1 channels primarily conduct Na^+^ and Ca^2+^ leading to a membrane depolarisation toward the action potential threshold [14].

Structurally, functional TRP channels are tetramers composed of four subunits, each comprising six transmembrane helices (TM1-TM6), with the loop between the 5^th^ and 6^th^ segments from all four subunits forming the channel pore. TRPA1 channels are characterised by an extended N-terminal region, containing several ankyrin repeat domains (ARD) consisting of pairs of antiparallel α-helices connected by β-hairpin motifs (14 repeats in the human channel). The N-terminal domain also contains EF-hand motifs that directly interact with calcium, allowing the channel to be activated by increased intracellular calcium levels. Conversely, deletions in the ARD region of the protein have been shown to impair protein trafficking to the plasma membrane [15]. Likewise, point mutations in the ARD region prevent calcium-dependent activation [16].

Exogenous and endogenous agonists bind to specific sites on the TRPA1 channel. For instance, electrophilic compounds, such as allyl isothiocyanate (AITC, the main component of the mustard oil), and cinnamaldehyde (found in cinnamon), react and form a covalent bond with nucleophilic amino acid residues, such as Cys (-SH) or Lys (-NH_2_), present in the cytoplasmic N-terminal domain in the channel [17, 18]. In contrast, non-electrophilic agonists such as menthol, THC, and carvacrol (a compound found in oregano) can activate the channel without covalent modification [19–21].

Importantly, calcium plays a complex modulatory role on channel activity. Raising extracellular calcium levels initially induces potentiation and subsequently inactivation by blocking the channel pore [22]. Furthermore, prolonged Ca^2+^-dependent activation results in channel desensitisation [23]. In addition, TRPA1 shows a weak voltage dependence, with the voltage-sensor like domain being located within an intracellular water-accessible crevice formed by transmembrane segments 1 to 4 [24]. Variants in this region have been associated to changes in the sensitivity of the channel to electrophilic compounds, voltage, extracellular calcium and membrane phosphoinositides [25].

Phosphatidylinositol-4,5-phosphate (PIP_2_) is a low-concentration anionic phospholipid component of the plasma membrane that has been shown to mediate the activity of several TRP channels, such as TRPV1 and TRM8. The role of PIP_2_ as a modulator of TRPA1 activity remains elusive. Intriguingly, some studies have shown that PIP_2_ inhibits TRPA1 [26], while others have argued for a role of PIP_2_ in its activation [27]. A previous study highlighted that PIP_2_ binding alters salt-bridge formation within the C-terminal domain of TRPA1, thereby regulating channel activity [27] while reductions in membrane PIP_2_ levels inhibit most mammalian TRP canonical channels (TRPC) [28, 29]. The S4-S5 linker region of TRPA1 appears to play a key role in channel gating and regulation, as it connects the voltage-sensing domain (S1-S4) to the pore domain (S5-S6) of the channel. In particular, the interaction between PIP_2_ and the S4–S5 linker may influence the flexibility of this region and, in turn, modulate the channel’s gating properties.

In this study, we investigated the impact of a rare variant on TRPA1 channel function and biophysical properties, in response to known TRPA1 agonists, following overexpression in both a heterologous system and primary sensory neurons. Our goal was to recapitulate the clinical phenotype and *in vitro* findings, to better understand the channel’s biophysical properties and explore potential implications for the future development of TRPA1-targeted therapeutics.

## Results

The novel TRPA1 variant M978V was identified in two members (father and daughter) of a family affected by painful sensory neuropathy. The father was assessed at the age of 59. His symptoms began in his late 30s with paraesthesia in the toes, which gradually spread to the level of his mid-calf, associated with numbness of the feet and foot pain, which he rated as 9/10 on a 10-point numerical rating scale. Foot pain was constant and not exacerbated by cold or exercise, and he denied muscle weakness. In addition to his sensory symptoms, he has had repeated episodes of infection of his toes and amputation of his second digit in both feet. He complained of urinary urgency but did not present any other autonomic symptoms. He used analgesia in the form of Co-codamol, tramadol, pregabalin and lamotrigine with only partial efficacy. His daughter also complained of painful feet (described in full detail below). He had an older son who had complained of some foot pain but had no evidence of neuropathy on clinical examination, and two unaffected sons. On examination, there was no limb weakness. Deep tendon reflexes were all present. Plantar reflexes were non-responsive. His gait was hesitant and Romberg’s sign was positive. On sensory examination, light touch was impaired to the forefoot, proprioception was impaired at the MTPs with small movements secure for the large movement. Vibration sense was absent to the knees, and pinprick sensibility was impaired to the mid-shin. Blood pressure was 170/90, and there was no postural drop.

Nerve conduction studies were consistent with a length-dependent sensory predominant axonal neuropathy. Genetic tests included a hereditary sensory neuropathy next-generation sequencing panel (including *ATL1*, *CCT5*, *DNMT1*, *FAM134B*, *NGF*, *NTRK1*, *RAB7A*, *SCN9A*, *SPTLC1*, *SPTLC2*, and *WNK1*). Further analysis using a pain channelopathy panel (*SCN9A*, *SCN10A*, *SCN11A* and *TRPA1*) revealed a missense variant in *TRPA1* present in heterozygosity: c.2932A>G. This was confirmed by Sanger sequencing. This results in a Methionine to Valine substitution at protein position 978 (p.M978V).

The daughter was assessed at the age of 28, at which time she had experienced bilateral pain on the underside of the heels for approximately a year. The pain was worse on waking and when she had been standing for a prolonged period. The pain had a burning quality, and typically she scored it as 6/10 on a numerical rating scale. She denied numbness but complained of paraesthesia in the toes. There was no weakness in the feet and no symptoms in the hands. There has been no change in her walking or balance. Occasionally, she would use simple analgesia such as NSAIDs or paracetamol. She did not have any autonomic symptoms. On examination, cranial nerves were normal. Tone, power and coordination of the limbs was normal. Deep tendon reflexes were all preserved, apart from ankle reflexes which were absent on the right and reduced on the left. On sensory examination, she complained of subjective reduction of pinprick up to the level of the mid foot bilaterally. Light touch, vibration sense and proprioception were all normal. Her gait was normal. Romberg’s sign was negative. Blood pressure was 120/80 lying with no postural drop. Screening blood tests did not show an underlying cause for neuropathy (full blood count, erythrocyte sedimentation rate, vitamin B12, folate, liver function test, bone profile, renal profile, plasma protein electrophoresis, thyroid function tests, and HbA1C were all normal or negative). Nerve conduction studies demonstrated normal sensory and motor conduction. A skin biopsy demonstrated a significantly reduced intra-epidermal nerve fibre density (4.2 fibres/mm, the lower limit of normal for her age and gender matched controls is 7.1 fibres/mm). These findings are consistent with a diagnosis of painful small fibre neuropathy. She was also heterozygous for the TRPA1 (M978V) variant.

The M978V missense variant is rare (allele frequency: 0.000121, 195/1613926 in the total population, 184/1179806 in the European (non-Finnish) population, according to GnomAD) and was identified in a family (the father being affected by painful neuropathy and the daughter by small fibre neuropathy). M978 is located in the proximal C-terminal region of the human TRPA1 channel (**Supplementary Figure 1A**), also known as “TRP box”, which is involved in the gating of the channel [30]. This region of the protein contains voltage-sensor domains (amino acids R975, K988 and K989 [31]) as well as the calmodulin binding site (AA 992-1008 [32]). Nearby, there are other binding sites for membrane lipids, including cholesterol, hepoxilin A3, and diacylglycerol, which can activate TRPA1 and mediate inflammatory mechanisms [33–36]. In addition, previous studies have shown that AA substitutions in this region affect the steady-state outward rectification and increase voltage sensitivity, while reducing sensitivity to AITC, such as K989A [31].

Although the amino-acid exchange M>V shows a relatively low Grantham score (21) [37], Methionine 978 is highly conserved in different mammalian species (**Supplementary Figure 1B**). Additionally, *in silico* prediction algorithms (such as Polyphen-2 [http://genetics.bwh.harvard.edu/pph2/] and SIFT [https://siftdna.org/index.html]) suggest that an AA exchange in this region of the protein is likely to affect the channel properties (Polyphen2: probably-damaging; SIFT: Deleterious). We thus sought to investigate the electrophysiological impact of the variant on the biophysical properties of TRPA1.

In a heterologous expression system, we assessed the current-voltage (I-V) relationship by applying a voltage-step protocol with the patch-clamp method (whole-cell configuration). To study the impact of the variant on TRPA1 activation and deactivation in control condition or in response to a TRAP1 agonist, we measured the channel’s voltage sensitivity and half-maximal activation potential. Current densities were assessed in WT and M978V, before and after application of AITC. Before AITC application (control condition), outward currents extrapolated at positive membrane potential (+100mV) were significantly increased in the presence of the M978V variant compared to its WT counterpart (WT: 20.02 pA/pF ± 2.59; M978V: 34.44 pA/pF ± 3.43, **Figure 1B**). On the other hand, inward currents (extrapolated at −100mV) remained largely unaltered (WT: −8.76 pA/pF ± 1.15; M978V: −9.82 pA/pF ± 0.97 S.E.M) (**Figure 1D**). We also assessed voltage sensitivity and gating properties in the presence of the novel variant. In control conditions, voltage sensitivity appeared unaltered (slope k: 44.14 ± 1.39 mV and 44.88 ± 2.65 for WT and mutant, respectively). Similarly, V_50_, extrapolated from voltage activation curves of WT and mutant, revealed no change (WT: 59.49mV ± 4.20; M978V: 50.95mV ± 7.61) (**Figure 1C**) as well as conductance-voltage relationship remained unaltered **(Figure 1E**). The reversal potential was also unchanged, suggesting that ion permeability is not altered when the variant is expressed (17.84 ± 4.86 mV and 18.40 ± 3.20 for WT and M978V, respectively, **Figure 1A**). Interestingly, reversal potentials between 0 and +10mV have been previously reported for the TRPA1 channel [38]; however, values closer to +20mV have also been observed, largely depending on the experimental conditions, particularly when the extracellular Ca^2+^ concentration is high [39]. This suggested that the presence of the variant primarily impacts the outward rectification of the channel prior to AITC application.

**Figure 1.**
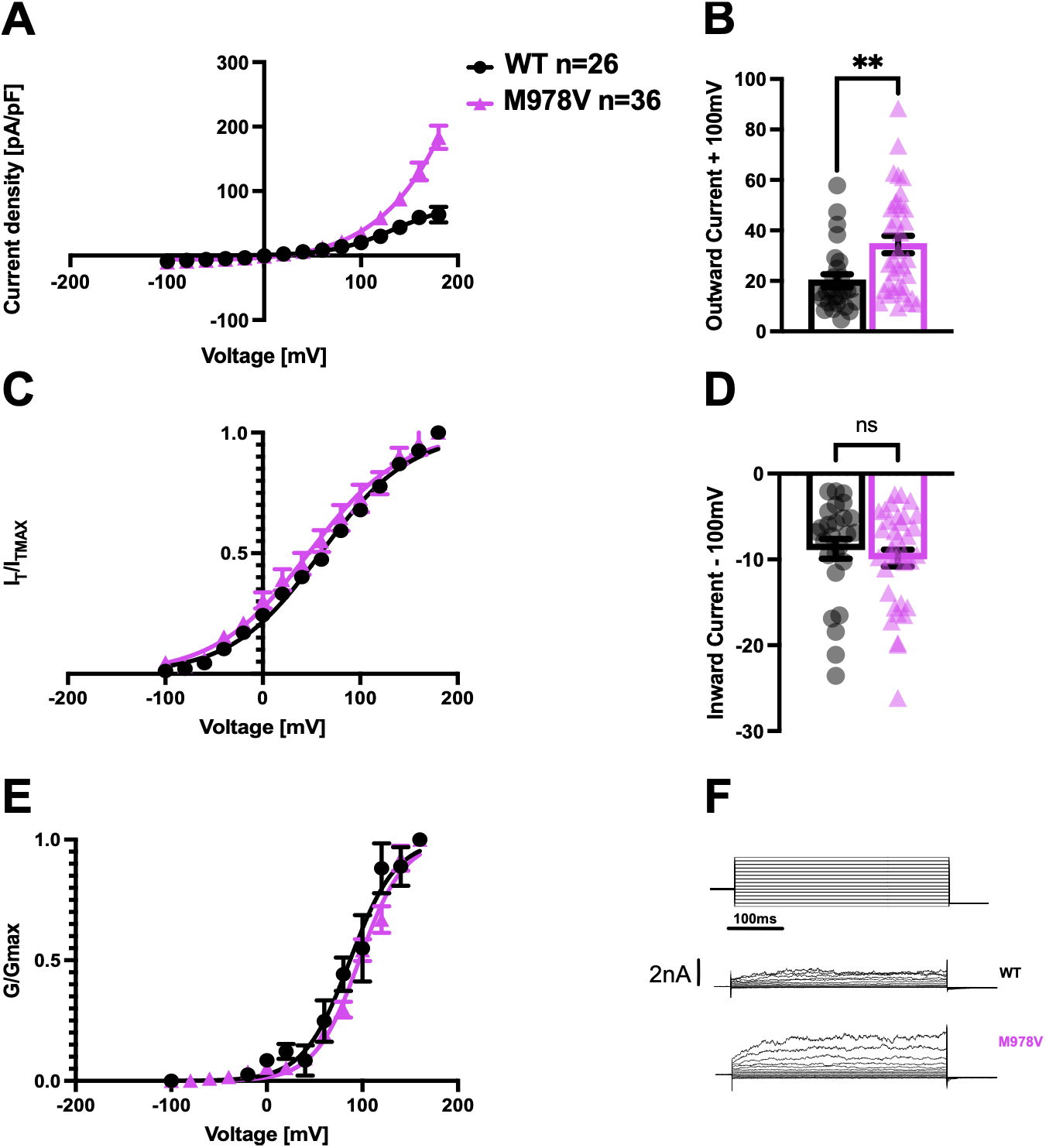
M978V increases outward current density of the human TRPA1 channel. Legend: (**A**) Average current-voltage (I-V) curve of hTRPA1-WT and hTRPA1-M978V in control conditions. Current density (pA/pF) was normalised by the cell membrane capacitance. Outward and inward currents, extrapolated at +100mV and −100mV, are shown in graph plots **B** and **D** respectively. (**C**) Mean steady-state activation curves were extrapolated from tail currents analysis at −70mV for both WT and mutant. (**E**) Average conductance-voltage (G-V) relationship was extrapolated from I-V curves, and G/G_max_ is normalized conductance. Solid lines are best fits to Boltzmann sigmoidal functions. Representative whole-cell current traces of HEK293T cells expressing hTRPA1-WT or hTRPA1-M978V in response to a two steps-voltage protocol (first step from −100mV to + 100mV, second step −70mV) are shown in (**F**).

We then sought to investigate the effect of the M978V variant on the channel activation and voltage sensitivity after application of AITC. In the presence of the agonist, we observed a left-ward shift in the half-maximal activation potential for the mutant channel (51.16 ± 4.05 mV and 26.81 ± 8.58 for WT and M978V, respectively, **Figure 2C**). This effect appears consistent with the position of the variant, which is located nearby one of the channel voltage sensors. The slope factor (k) was also shifted towards less depolarised potential, indicating that the voltage sensitivity is also affected by the variant (WT: 47.32mV ± 1.75; M978V: 35.15mV ± 2.47). Outward currents also appeared significantly increased (WT: 27.29 pA/pF ± 3.99; M978V: 148.2 pA/pF ± 20.86, **Figure 2B**), as well as inward currents (WT: −7.35 pA/pF ± 1.49; M978V: −13.80 pA/pF ± 1.95, **Figure 2D**). Notably, for the mutant channel the current decays at strongly depolarised potentials (**Figure 2A**), a phenomenon that has been previously observed and reported as voltage-dependent inactivation [40].

**Figure 2.**
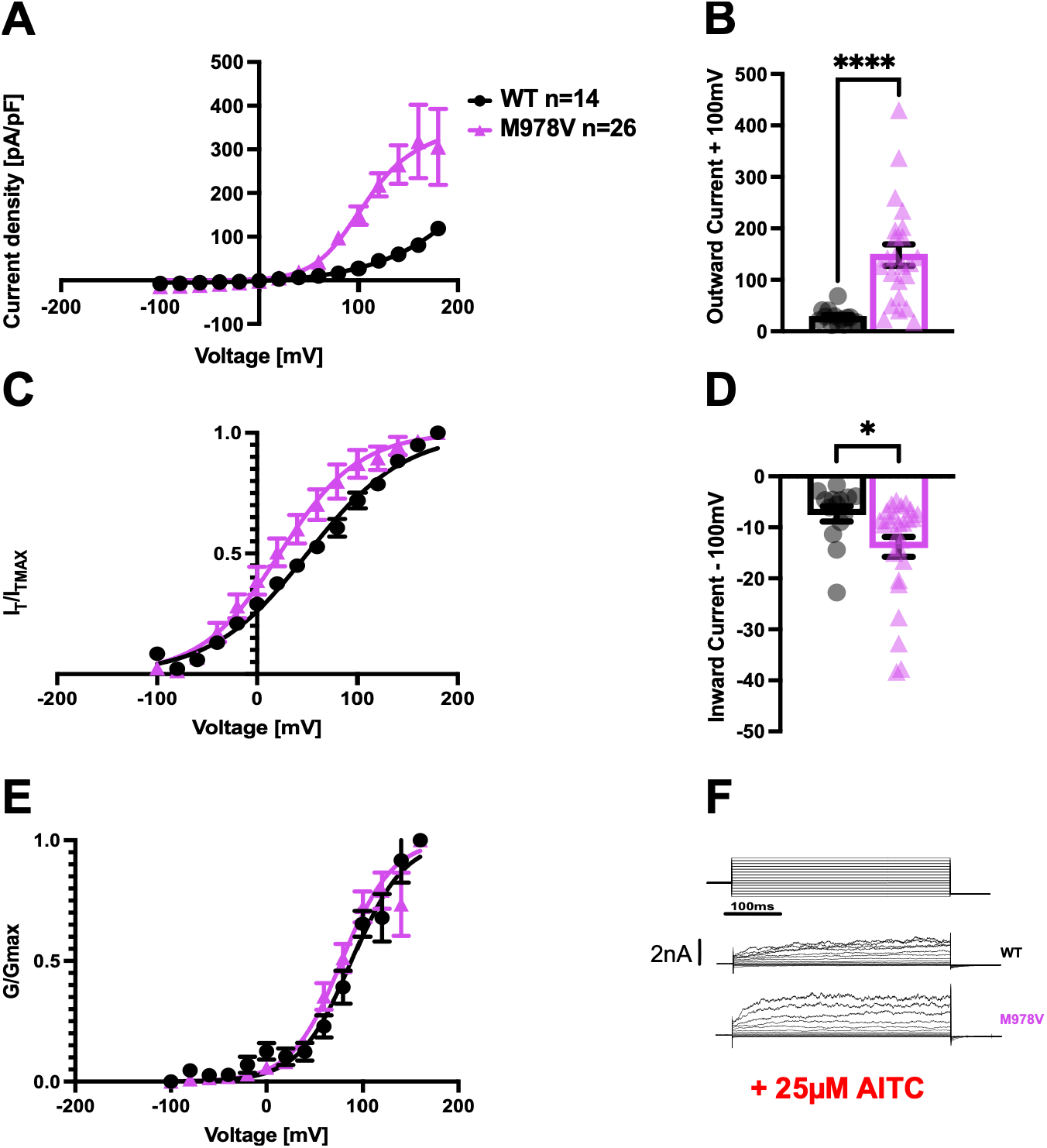
M978V confers gain-of-function properties to the human TRPA1 channel after 25µM AITC application. Legend: (**A**) Average I-V curve of hTRPA1-WT and hTRPA1-M978V in after AITC application. Current density (pA/pF) was normalised by the cell membrane capacitance. Mean steady-state activation curves were extrapolated from tail currents analysis at −70mV for both WT and mutant after AITC application (**B**). Outward and inward currents, extrapolated at +100mV and −100mV, are shown in **C** and **D**, respectively. Average conductance-voltage relationship was extrapolated from I-V curves obtained from voltage step protocols, and G/G_max_ is normalized conductance (**F**). Solid lines are best fits to Boltzmann sigmoidal functions. Representative whole-cell current traces of HEK293T cells expressing hTRPA1-WT or hTRPA1-M978V in response to the indicated voltage-step protocol are shown in (**E**) after perfusion with 25µM AITC.

In these experimental conditions, however, the reversal potential appeared unaltered (0.4 ± 6.36 mV and 7.5 ± 4.41 for WT and mutant, respectively, **Table 1**), arguing against an altered ion permeability in the M978V channel, after perfusion with AITC. Importantly, the G/G_max_ curve (channel conductance) measured at the steady state and extrapolated from the I-V curve was not altered in WT versus mutant condition, either in control conditions (showing a V_50_ of 89.58 ± 11.83 mV and 100.2 ± 6.84 mV for WT and M978V, respectively, p = 0.69, **Figure 1E**) or after stimulation with 25µM AITC (reporting a V_50_ of 95.32 ± 6.28 mV for WT and 93.93 ± 4.35 mV for M978V, p = 0.85, **Figure 2E**). This suggests that, although voltage sensitivity is altered in the presence of the variant, with a significant left-ward shift of the V_50_, once the mutant channel is open, the conductance of the channel remains largely unaltered. M978V exhibited increased voltage sensitivity under both control and AITC-stimulated conditions. This effect was more pronounced after AITC application, with the variant shifting the V_50_ toward less depolarised potentials and altering the channel’s open probability. The main electrophysiological parameters are summarised in **Table 1**.

**Table 1.** Summary of electrophysiological parameters of WT and M978V TRPA1 biophysical properties.

| Condition | $V_{50}$ (mV) | Slope (mV) | $E_{rev}$ (mV) | Ratio<br>(inward/outward current<br>pA/pF) | Protocol |
| --- | --- | --- | --- | --- | --- |
| Control<br>(ECS with $Ca^{2+}$) | WT = 59.49 ± 4.2<br>M978V = 50.95 ± 7.6<br>$P=0.309$<br>two-tailed t test | WT = 44.14 ± 1.4<br>M978V = 44.88 ± 2.6<br>$P=0.796$<br>two-tailed t test | WT = 17.84 ± 4.8<br>M978V = 18.40 ± 3.2<br>$P=0.92$<br>two-tailed t test | WT = -0.51 ± 0.06<br>M978V = -0.32 ± 0.03*<br>$P=0.012$<br>two-tailed t test | Voltage-Step |
| AITC 25μM<br>(ECS with $Ca^{2+}$) | WT = 51.16 ± 4.0<br>M978V = 26.81 ± 8.5*<br>$P=0.0341$<br>two-tailed t test | WT = 47.32 ± 1.7<br>M978V = 35.15 ± 2.4**<br>$P=0.0014$<br>two-tailed t test | WT = 0.40 ± 6.3<br>M978V = 7.56 ± 4.4<br>$P=0.59$<br>two-tailed t test | WT = -0.26 ± 0.03<br>M978V = -0.13 ± 0.03*<br>$P=0.0126$<br>two-tailed t test | Voltage-Step |
| Control<br>(No $Ca^{2+}$)<br>(No PIP<sub>2</sub>) | WT = 27.66 ± 6.3<br>M978V = 36.27 ± 7.2<br>$P=0.375$<br>two-tailed t test | WT = 45.79 ± 2.7<br>M978V = 54.94 ± 4.8<br>$P=0.097$<br>two-tailed t test | WT = -11.48 ± 7.2<br>M978V = -10.52 ± 6.7<br>$P=0.92$<br>two-tailed t test | WT = -0.38 ± 0.06<br>M978V = -0.48 ± 0.04<br>$P=0.181$<br>two-tailed t test | Voltage-Step |
| AITC 25μM<br>(No $Ca^{2+}$)<br>(No PIP<sub>2</sub>) | WT = 20.22 ± 9.1<br>M978V = 29.27 ± 7.1<br>$P=0.44$<br>two-tailed t test | WT = 46.47 ± 4.0<br>M978V = 54.64 ± 6.7<br>$P=0.311$<br>two-tailed t test | WT = -18.55 ± 7.0<br>M978V = 4.49 ± 4.6**<br>$P=0.0087$<br>two-tailed t test | WT = -0.33 ± 0.07<br>M978V = -0.37 ± 0.05<br>$P=0.611$<br>two-tailed t test | Voltage-Step |

**Legend:** $V_{50}$ and slope values were calculated from Boltzmann fits of tail current amplitudes measured at -70mV, when tail amplitude is proportional to the channel open probability and the driving force is constant. The reversal potential ( $E_{rev}$ ) was determined by steady-state linear interpolation of the $I-V$ relationship near the zero-current region, with the x-intercept representing $E_{rev}$ .

TRPA1 can be bimodally activated by calcium, in that calcium, at low concentrations, potentiates the channel activation (e.g., in response to electrophile agonists such as AITC), while at higher concentrations it leads to channel desensitisation. It has been shown that AITC *per se* can trigger the release of intracellular calcium [41]. Given that the gating behaviour of the TRPA1 channel is closely dependent on the presence of external/internal Ca^2+^, as previously reported [13], we assessed current density in nominally Ca^2+^-free extracellular solution (**Supplementary Figure 1**). A voltage-ramp protocol (from –100 mV to +100 mV over 500 ms) was applied to assess the activation properties of the channel (I-V curve) over time, in the presence of 25µM and 100µM AITC (**Supplementary Figure 1A and C**, respectively. The voltage-ramp protocol allows the gradual depolarisation of TRPA1 to reduce channel desensitisation. In both WT and mutant conditions, we observed a linearisation of the I-V curve, which appeared more pronounced at the highest AITC concentration (100µM).

Previous studies have shown that AITC can induce exocytosis and facilitate membrane expression of the human TRPA1 channel, as suggested by measures of membrane capacitance [42]. AITC can modulate membrane channel trafficking by recruiting TRPA1-containing vesicles to the membrane and activating TRPA1 channels on the ER, promoting further release of calcium from intracellular stores [43]. AITC also induces releases of substance P and CGRP [7, 44], neuropeptides which are secreted during inflammation. Surface expression of the channels has also been linked to changes in temperatures, such as warm and cold noxious stimuli [8]. Since the mutant channel showed higher current density upon transient over-expression in a heterologous system, especially after AITC stimulation, we examined whether this effect involved altered channel membrane expression.

Cells were either incubated with AITC or vehicle for different incubation times (5 min and 30min) in both absence and presence of extracellular calcium. We then quantified the surface expression of WT or mutant channel as a ratio of the mean fluorescence intensity of the plasma membrane compared to that of the cytoplasm, using an antibody targeting TRPA1. Our data show that after incubation with AITC for 5 or 30 min, the surface expression of the hTRPA1 M978V channel is increased, compared to WT, in nominally Ca^2+^-free bath solution, with a more prominent effect at longer incubation times (**Figure 3B** and **3D**; two-way ANOVA multiple comparison, 5 min incubation: p value 0.3597 ns, p value 0.0045 **, p value < 0.0001 ****; 30 min incubation: p value 0.01 *, p value 0.0033 **, p value 0.0001 ****).

**Figure 3.**
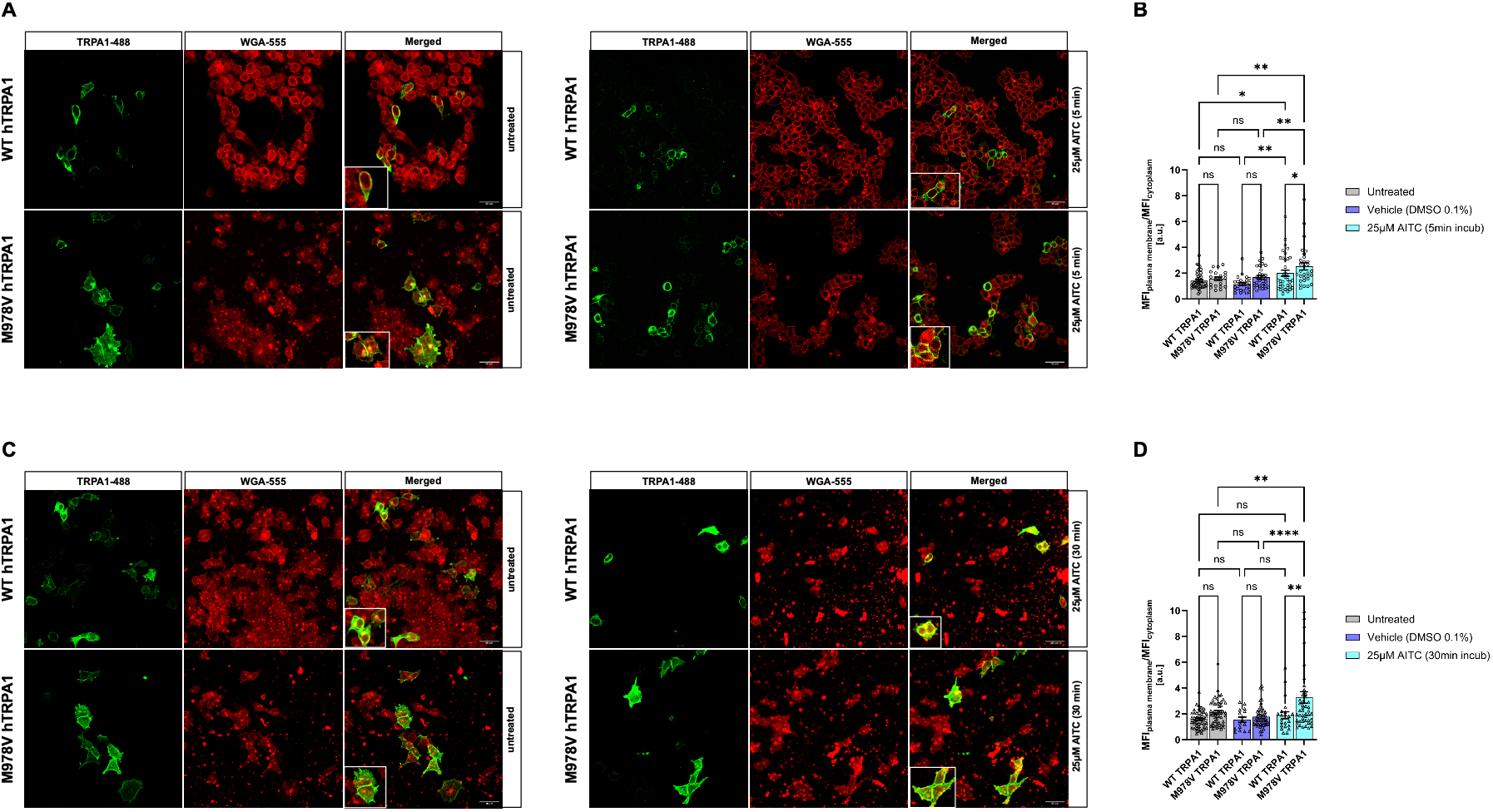
Surface membrane expression is enhanced in the presence of the TRPA1 M978V variant. Legend: Representative images of HEK293T cells transiently transfected with either WT or mutant channel are shown in **A** and **C**; cells were counter-stained with a TRPA1 antibody (shown in green). Wheat germ agglutinin (WGA) antibody was used to visualise the cell membrane (shown in red). Merged images show the overlap between WGA-stained plasma membranes and TRPA1-positive cells. Panel **B** shows the quantification of TRPA1 plasma membrane expression, expressed as the plasma membrane mean fluorescence intensity (MFI) to cytoplasmic MFI ratio, following a 5-min incubation with either Ca²⁺-free extracellular solution (untreated, WT n = 40 and M978V n = 20), vehicle (0.1% DMSO, WT n = 25 and M978V n = 30), or 25 µM AITC (WT n = 35 and M978V n = 36) in the absence of extracellular Ca²⁺. Panel **D** reports MFI after 30 min incubation with Ca²⁺-free extracellular solution (WT n = 46 and M978V n = 50), vehicle (WT n = 16 and M978V n = 36) or 25µM AITC (WT n = 50 and M978V n = 47). Scale bar: 40µm. All grouped data are mean ± SEM,* p < 0.05, ** p < 0.01, *** p < 0.001; **** p < 0.0001.

In contrast, in the presence of Ca^2+^_Ext_, shorter incubation (5 min) with 25µM of AITC did not seem to significantly increase the surface (relative to cytoplasmic) expression of the mutant channel compared to WT, although it did show increased mean fluorescence intensity, on the plasma membrane, compared to control conditions (untreated and vehicle-treated cells, **Supplementary Figure 2 upper panel B**). After longer exposure to either ECS, DMSO or AITC (30 min), the surface expression of the mutant channel appeared increased compared to that of the WT channel in all three treatment conditions; however, there was no relative additional increase in the presence of the variant upon AITC treatment compared to its untreated state (**Supplementary Figure 2 lower panel B;** two-way ANOVA multiple comparison, 5 min incubation: p value 0.4139 ns, p value 0.8197 ns, p value < 0.0001 ****; 30 min incubation: p value 0.3708 ns, p value 0.965 ns, p value 0.0001 ****).

Given the proximity of M978 to a series of basic residues on the human TRPA1 structure model, we then hypothesised that the variant may alter the binding affinity of the channel for PIP_2_. To test our initial hypothesis, we conducted 5×10μs coarse-grained molecular dynamics simulations of the hTRPA1 channel in a POPC bilayer containing 10% PIP_2_ (**Figure 4A**). Using VolMap analyses, we showed that PIP_2_ binds at the basic region, primarily between residues 800 and 900 near the M978 variant, in agreement with our initial hypothesis (**Figure 4B** and **4C**). This allowed us to pick the PIP_2_ binding site nearest to M978 and convert our coarse-grained representation to an all-atom representation. The simulations were then conducted for a further 500 ns with 5 repeats. Our simulations highlighted that the predicted PIP_2_ binding site comprised of N855, L871, R872, Y874, V875, I878 and R975 is located on the S4-S5 helices (**Figure 4D** and **4E**). We also assessed the stability of the PIP_2_ headgroup in the binding pocket and showed that the PIP_2_ headgroup remains stable throughout 500 ns of simulations, validating the nature of the binding pocket (**Figure 4F**).

**Figure 4.**
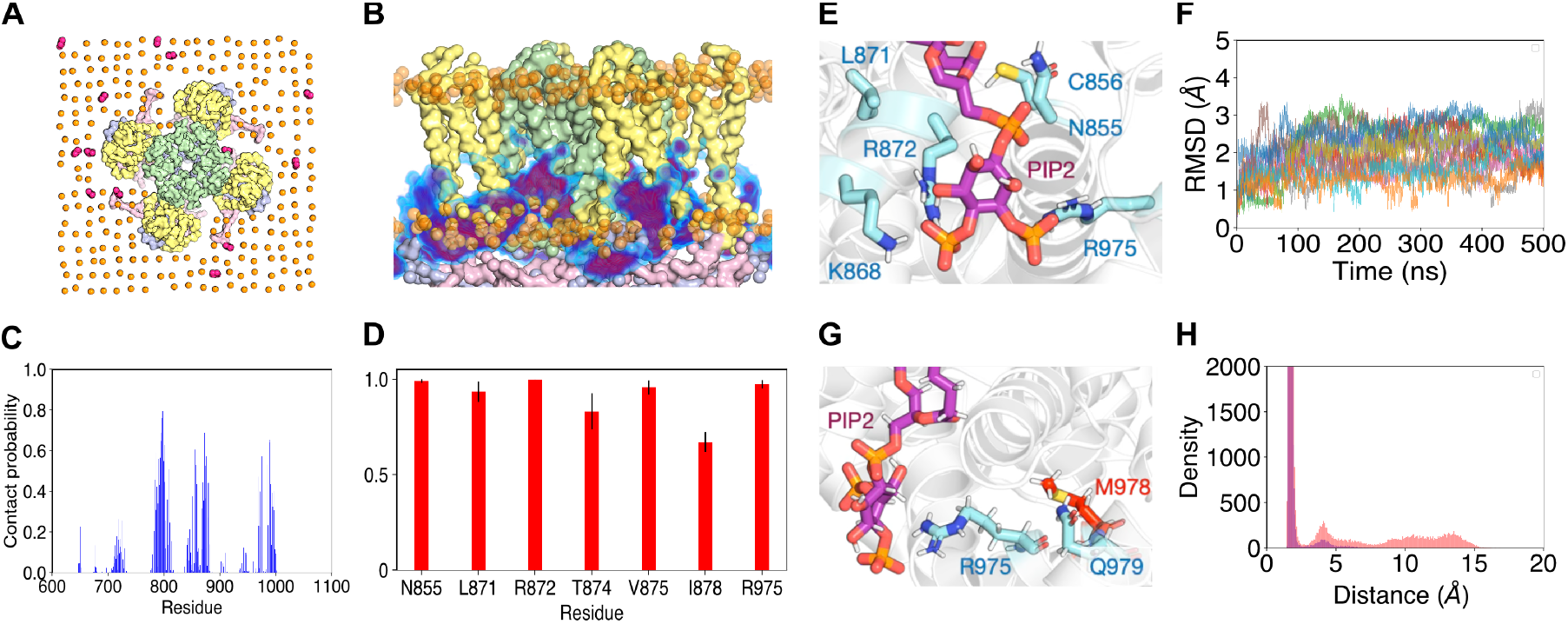
MD simulations reveal a novel binding site for PIP2 in the human TRPA1 channel. Legend: Panel **A** depicts a POPC bilayer containing 10% PIP_2_ as a substrate for the human TRPA1 channel. **B** and **C** show the structural model of the PIP_2_ binding site, between residues 800-900 of the channel, Representations of the residues involved in the binding site are shown in **D** and **E**. Panel **F** shows the stability of PIP_2_ in the binding pocket over the course of the MS simulations (500ns). Panel **G** and **H** indicate the position of the M978V residue. All simulations were carried out using the Martini2.2 biomolecular forcefield.

We then asked whether introducing the variant of interest, M978V, would influence the interaction of R975 to PIP_2_, given the proximity to Q979 and R975 residues (**Figure 4G**). Thus, we conducted MD simulations with M978V mutant for 500 ns x 5 repeats. Here, we showed that the variant influences the distance between R975 and the PIP_2_ headgroup (**Figure 4H**), suggesting that the variant can impact the interaction between the R975 residue and the PIP_2_ binding site. C-alpha atom root-mean-square fluctuation (RMSF) of human TRPA1 did not reveal any significant impact of the mutated aminoacid on the protein flexibility and stability (**Supplementary Figure 4A**).

Overall, these predictions suggest that the M978V variant is likely to allosterically disrupt the nature of the PIP_2_ binding site on the TRPA1 channel.

Our MD simulations (**Figure 4G** and **4H**) revealed the proximity of the M978V variant to one of the PIP_2_ binding sites; we thus sought to experimentally investigate the effect of the PIP_2_, in particular, whether the gain-of-function properties of TRPA1 caused by M978V in response to AITC depend on the presence of PIP_2_. To do so, we co-transfected either WT or M978V TRPA1 together with *Danio Rerio* voltage-sensitive phosphatase Dr-VSP, an enzyme which degrades PIP_2_ when membrane potential exceeds +50 mV [45, 46]. We employed the patch clamp technique and the two step-voltage protocol previously applied (**Figure 5E**).

**Figure 5.**
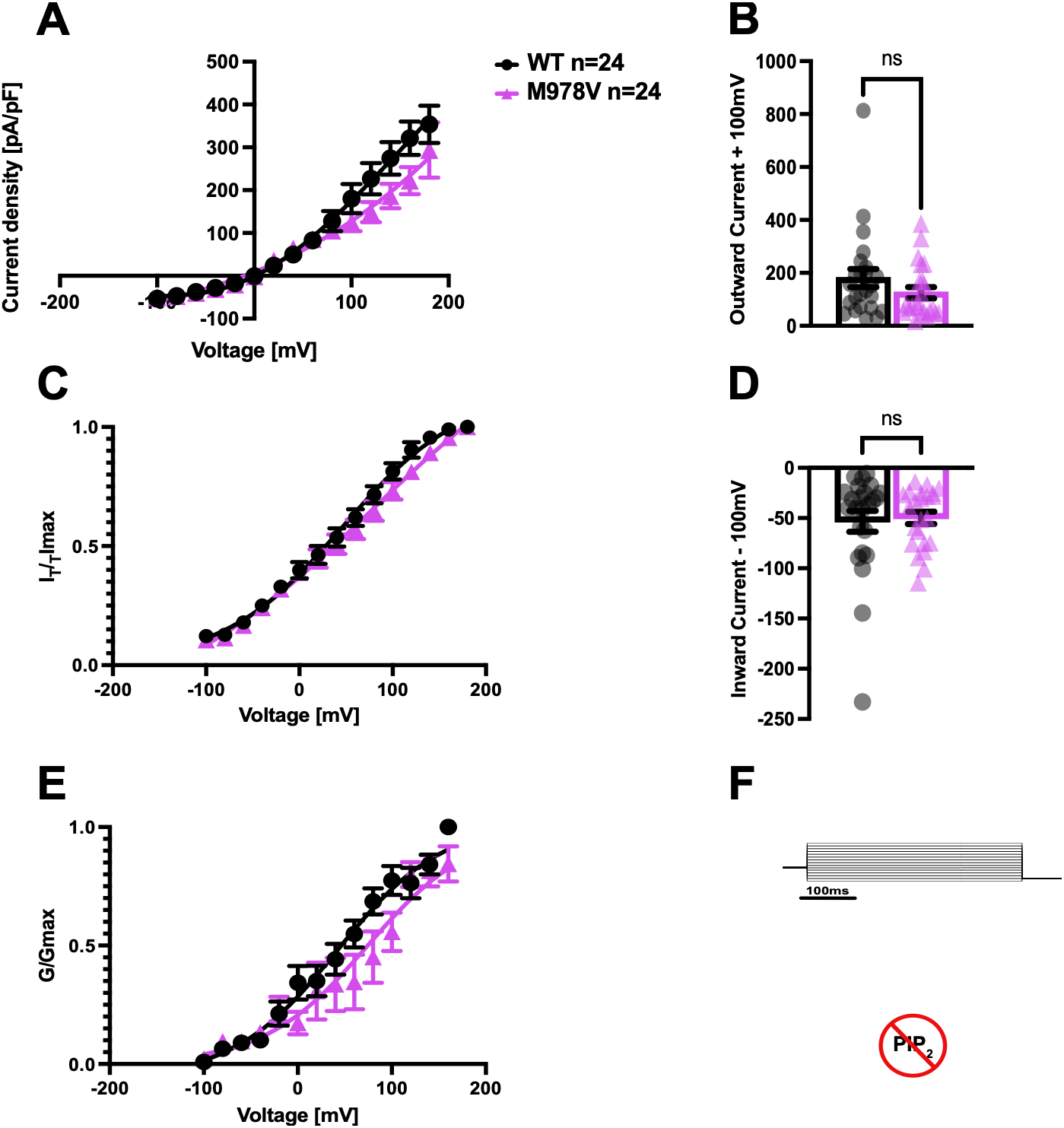
Depletion of PIP_2_ increases current density for both WT and mutant channel in control conditions. Legend: Panel **A** shows I-V curve derived from current densities of hTRPA1-WT and hTRPA1-M978V in control conditions. Current density (pA/pF) was measured in WT and M978V channels, in presence of the DrVSP (voltage-sensitive phosphatase), and normalised by the cell membrane capacitance. Mean steady-state activation curves were extrapolated from tail currents measured at −70mV for both WT and mutant (**C**). Panel **B** depicts outward (at +100mV) and inward (at −100mV) currents (upper and lower panel respectively). Panel **D** shows average G-V relationship was extrapolated from I-V curves obtained from voltage-step protocols. Normalised conductance (G/G_max_) was plotted as a function of test potential and fitted with a Boltzmann function. Solid lines are best fits to Boltzmann sigmoidal functions. Representative whole-cell current traces of HEK293T cells expressing hTRPA1-WT or hTRPA1-M978V in response to the indicated voltage-step protocol are shown in (**E**).

Our *in vitro* data (**Figure 5A**) show that, in the absence of extracellular calcium, depletion of intracellular PIP_2_ via DrVSP increased current densities for both WT and M978V channels compared to when PIP_2_ is endogenously expressed (**Figure 1A**), particularly at positive potentials (> +50mV). However, in basal control conditions open probability (V_50_) and voltage sensitivity (slope K factor and G/G_max_) when comparing WT and mutant were largely unaffected (**Figures 5C** and **E**). Tail current analysis revealed that V_50_ was 27.66 ± 6.37 mV and 36.27 ± 7.26 mV for WT and mutant, respectively, with the slope (k factor) being 45.79 ± 2.71 (WT) and 54.94 ± 4.88 (M978V) (**Figure 5C**). Consistently, after applying a Boltzmann fit to the G-V curve, conductance was not significantly changed between the two genotypes (WT=39.88 ± 13.82mV, M978V=68.70 ± 20.52mV, p value=0.262, two-tailed t test, **Figure 5E**). However, both V_50_ and reversal potential (E_rev_) where affected when comparing basal condition in the presence and absence of PIP_2_ (**Table 1**). As previously shown [25], in the absence of external calcium, the inner cavity of the sensor domains (S1-S4 linkers) is normally occupied by endogenous PIP_2_, ensuring proper channel gating. Depletion of PIP_2_ (for instance *via* PLC activation) has been shown to sensitise TRPA1 and facilitates its activation, enhancing its response to electrophilic compounds like AITC [26]. Our results are in line with these findings, and show that depletion of PIP_2_ indeed leads to increased current density for the hTRPA1 channel. Interestingly, under these experimental conditions, we did not observe any significant difference between WT and mutant channel, with I-V, V_50_ and G-V largely unaltered (**Figure 5A, 5C** and **5D**).

Depletion of PIP_2_ has been shown to sensitise TRPA1 and facilitates its activation, enhancing its response to electrophilic compounds like AITC [26]. We thus sought to investigate the impact of the M978V variant after depletion of PIP_2_ and in the presence of AITC. Both outward and inward currents were increased in WT and M978V channels compared with control conditions, with a more prominent effect for the WT channel. Specifically, outward currents (extrapolated at +100mV) were 463.7 ± 76.74 pA/pF and 146.5 ± 28.11 pA/pF respectively) while inward currents (extrapolated at −100mV) were −115.6 ± 29.18 pA/pF and −43.44 ± 10.31 pA/pF for WT and mutant, respectively (**Figures 6B** and **D**). Consistently, also channel conductance appeared higher in WT (**Figure 6E**), whereas open probability, reversal potential (E_rev_) and intrinsic voltage-dependent gating, extrapolated from tail current amplitudes, remained largely unchanged (**Figure 6C**) (V_50_ was 20.22 ± 9.16 mV and 29.27 ± 7.19 mV, the slope factor was 46.47 ± 4.00 mV and 54.64 ± 6.71 mV and the reversal potential was −18.55 ± 7.0mV and 4.49 ± 4.6 mV for WT and M978V, respectively, see **Table 1**). Although the voltage dependence of activation (V₅₀) derived from tail current amplitudes was unchanged, the V₅₀ obtained from conductance–voltage (G–V) relationship displayed a leftward depolarising shift for the WT channel (28.61 ± 6.48mV for WT and 97.62 ± 15.23mV for M978V, P=0.0002, two tailed t test, see **Table 1**). Importantly, the gain-of-function phenotype previously associated with the M978V variant was no longer observed. In conclusion, depletion of PIP_2_ in the presence of AITC preferentially affected WT channels over mutant channels, as reflected by larger current densities, higher conductance, and a leftward shift in the voltage dependence of half-maximal activation. This differential response abolished, and indeed reversed, the gain-of-function phenotype previously associated with the M978V variant.

**Figure 6.**
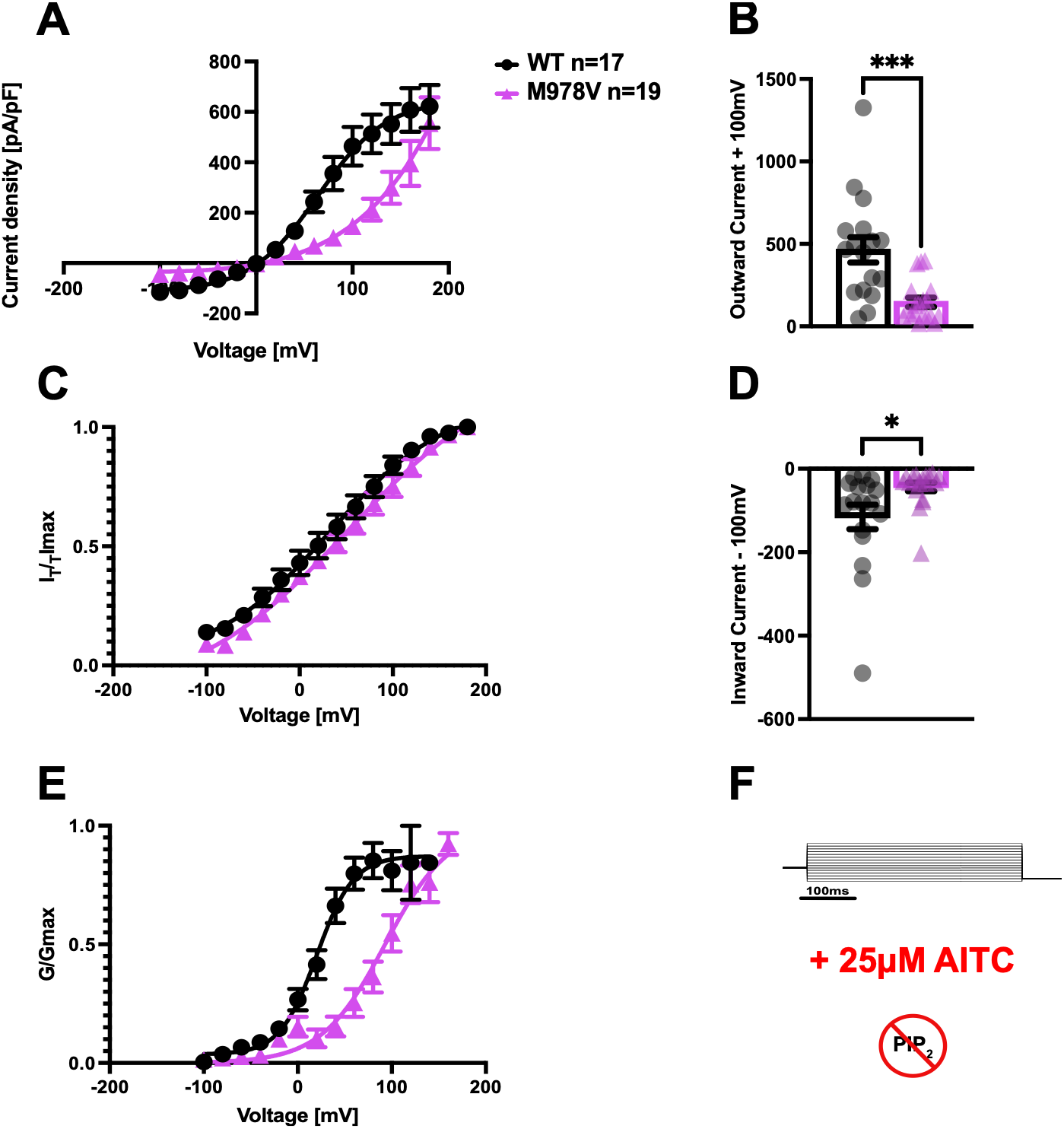
Depletion of PIP_2_ abolishes the GOF effect of M978V in response to AITC. Legend: I-V curve calculated from current densities of WT and mutant TRPA, in presence of the DrVSP, and normalised by the cell membrane capacitance are shown in panel **A**. Tail currents were extrapolated at −70mV (**B**). Panel **C** and **D** depict outward (at +100mV) and inward (at −100mV) currents, respectively. Averaged G-V relationship was extrapolated from I-V curves obtained from voltage-step protocols. Normalised conductance (G/G_max_) was plotted as a function of test potential and fitted with a Boltzmann function (**F**). Solid lines represent best fits to Boltzmann sigmoidal functions. The voltage-step protocol applied and representative whole-cell current traces of HEK293T cells expressing hTRPA1-WT or hTRPA1-M978V in response to the indicated voltage-step protocol are shown in (**E**).

## Discussion

This study demonstrates that M978V, a rare *TRPA1* genetic variant identified in a father and daughter with painful sensory neuropathy, is a gain-of-function (GOF) variant that alters PIP_2_-dependent action by the electrophilic agonist AITC. A few *TRPA1* variants have previously been linked to pain. For instance, the N855S was the first clinically relevant variant to be described in a large Colombian pedigree with familial episodic pain syndrome (FEPS). FEPS is an autosomal dominant disorder characterised by episodes of severe upper body pain, triggered by physical stress, fasting, or cold. Sequencing of candidate genes identified a missense AA change (N855S) in the S4 of the human *TRPA*1 channel resulting in a GOF following agonist stimulation (more than fivefold larger inward current at normal resting potentials compared with the wild-type channel [13, 47]). A second pedigree with severe episodic truncal pain was subsequently found to carry A172V, an ankyrin-domain variant that likewise enhanced AITC-evoked currents [48]. Importantly, both N855S and A172V variants manifested as pain channelopathies without clinical evidence of an associated neuropathy. More recently, the genetic basis of neuropathic pain has been increasingly investigated [49]. Advances in next-generation sequencing and gene-burden analyses have strengthened the association between rare *TRPA1* variants and painful neuropathies and other pain disorders, including fibromyalgia [50–52]. Yet, most of these variants remain functionally uncharacterised and are supported only *by in silico* pathogenicity predictions.

Calcium plays a complex, bidirectional role in regulating TRPA1 activity. Both intracellular and extracellular calcium can activate TRPAl, potentiating its response to several electrophilic and non-electrophilic agonists, while sustained calcium influx subsequently promotes prolonged channel inactivation [41, 44, 53, 54]. For this reason, we sought to investigate the gating properties of the M978V mutant channel in the presence and absence of extracellular calcium.

Our results show that in HEK293T cells, in the presence of extracellular calcium, both the WT and mutant channels exhibit weak voltage dependence with characteristic outward rectification, consistent with previous reports [31, 38], with outward currents enhanced in the presence of the variant (**Figure 1**). Under basal conditions, reversal potential (E_rev_) as well as open probability (V_50_) remained unaltered, suggesting that the variant does not affects ion permeability or intrinsic gating at rest.

However, AITC stimulation of the M978V variant resulted in increased current density at both positive and negative membrane potentials, an increased inward-to-outward current ratio, and shifts in both the half-activation voltage (V₅₀) and voltage sensitivity (slope factor *k*) derived from tail currents (**Figure 2**). Given that neither the reversal potential nor the conductance were altered, we propose that the effect of the variant is localised to the voltage-sensor-like domain (S4–S6 linker) rather than the pore domain, which determines ion permeability and ion conductance.

Given the complex role of Calcium, we next investigated whether the gain-of-function phenotype extended beyond channel gating. In addition to enhancing channel activity, AITC stimulation increased expression of the M978V mutant on the plasma membrane in absence of extracellular calcium, particularly after prolonged AITC stimulation (**Figure 3**). However, channel expression was not increased in the mutant relative to WT following 5 min of AITC treatment (**Supplementary Figure 2B**). Although mutant surface expression was higher than WT at longer incubation times, no differences were observed between untreated, vehicle, and AITC-treated conditions. As previously reported, prolonged AITC exposure can induce TRPA1 desensitisation [55]. Extracellular calcium may also contribute to channel desensitisation, potentially affecting the WT channel more than M978V, consistent with the higher basal current density observed for the mutant under control conditions. Interestingly, in the absence of both intracellular and extracellular calcium, we observed an increasingly pronounced linearisation of the I-V relationship, for both WT and mutant channels (**Supplementary Figure 1B and D**). This reduced channel rectification was more pronounced in the presence of the M978V variant, suggesting altered gating transitions.

Phosphatidylinositol 4,5-bisphosphate (PIP_2_) is a lipid cofactor that regulates numerous ion channels through electrostatic interactions with basic residues within their cytoplasmic domains [56]. However, its action on human TRPA1 remains elusive. Several studies have reported that PIP_2_ positively regulates TRPA1 [57], enhancing channel activity in inside-out patch recordings [36]. More recently, PIP_2_ has been proposed to stabilise the open state of the channel, thus potentiating its activation [25]. In contrast, other studies have shown that PIP_2_ down-regulates TRPA1 [26], reporting that PIP₂ depletion facilitates channel activation in inside-out patches [58], consistent with an inhibitory role for this phospholipid [26]. Finally, others report no effect [59].

Our *in vitro* data show that DrVSP-mediated hydrolysis of PIP_2_ increased current density for both WT and mutant channels, in basal conditions (**Figure 5**) and after AITC stimulation (**Figure 6**). Critically, PIP_2_ depletion abolished the M978V gain-of-function effect; once PIP_2_ was hydrolysed, the mutant no longer displayed the activating effect. This provides a mechanistic link between the M978V variant and PIP_2_. PIP_2_ binds the S4–S5 linker, and AITC activates TRPA1 allosterically through covalent modification of cysteines (Cys621, Cys665) within the cytoplasmic N-terminal domain [60, 61]). Because M978V lies close to the proposed PIP_2_-binding site, we hypothesise that the variant enhances allosteric coupling between the ligand-sensing domains and the channel gate in a PIP_2_-dependent manner. PIP_2_ depletion might impair allosteric coupling between the AITC-binding site and the channel gate through the PIP_2_-binding site adjacent to M978V. In this model, PIP_2_ would facilitate AITC-dependent activation of the mutant channel rather than being a parallel modulator. This interpretation is further supported by the V₅₀ values, which are differentially affected by PIP₂ depletion depending on the method of measurement. Tail-current analysis reports the steady-state open probability and is largely independent of gating kinetics, whereas G–V curves are derived from macroscopic currents during depolarisation and are therefore shaped by both activation and inactivation. Only the G– V V₅₀ shifted after PIP_2_ depletion (**Figure 6**), thus suggesting that the variant is primarily affecting the gating kinetics rather than reflecting a real impact on the channel’s intrinsic voltage dependence.

Additionally, an important consideration is that the M978V variant is reported at a heterozygous GnomAD frequency of ∼1×10⁻⁴ suggesting that it is unlikely to be fully penetrant and is more appropriately viewed as a risk-conferring allele than a monogenic cause of disease. However, the drivers for channel activation *in vivo* (including the relative contribution of environmental versus endogenous stimuli) are not entirely known. We propose that the presence of the variant lowers the activation threshold of nociceptive afferents, thereby increasing susceptibility to pain in the presence of additional genetic, environmental, or physiological factors.

Relevant endogenous TRPA1 activators include reactive oxygen species (ROS), methylglyoxal (MG), and noxious heat and cold [4, 5, 62, 63], all of which can be elevated under pathological conditions. This may be particularly relevant in the distal extremities, such as the hands and feet, which are often among the first regions affected in peripheral neuropathies. Further studies are warranted to determine whether these rare TRPA1 variants exhibit altered sensitivity to ROS and MG. This question is especially pertinent in the context of diabetes and diabetic neuropathy, where MG accumulates as a consequence of hyperglycemia, while ROS production is increased due to chronic hyperglycemia, oxidative stress, and microvascular dysfunction. In our model, the gain-of-function phenotype is translated into a clinical manifestation only when combined with appropriate disease-promoting stimuli. Identifying these factors, and determining which endogenous or exogenous signals drive M978V-TRPA1 activation *in vivo*, will be essential for understanding the pathogenic mechanisms underlying this painful neuropathy.

Finally, our findings have important therapeutic implications. TRPA1 has already been actively pursued as an analgesic target [64]; however, relatively few small molecule TRPA1 antagonists have advanced to clinical trials, and we do not yet have unequivocal evidence of efficacy in patient populations [65, 66].

Our data demonstrate that the rare M978V gain-of-function phenotype is dependent on PIP_2_. In this new light, the protein–lipid interaction, rather than the channel pore itself, may represent a more appropriate therapeutic target. If M978V indeed remodels PIP₂-mediated allosteric coupling, then selectively modulating TRPA1–PIP₂ interactions could provide a novel mechanism-based therapeutic strategy for the treatment of TRPA1-mediated painful neuropathies. This would require stratifying patients by genotype, specifically identifying those subjects carrying rare TRPA1 variants modulated by PIP_2_, in line with the growing recognition that personalised approaches to pain management are needed [67].

## Materials and Methods

### Participants consent

All participants provided written informed consent in accordance with the Declaration of Helsinki.

### HEK293T cell culture and transfection

Human embryonic kidney cells (HEK293T) were grown in Dulbecco modified Eagle’s culture medium supplemented with GlutaMAX^TM^ (DMEM Gibco, 15440544, ThermoFisher Scientific, United Kingdom), containing 10% fetal bovine serum and maintained under standard conditions (37°C) in a humidified and controlled atmosphere (5% CO_2_). Cells were transiently transfected with the jetPEI transfection reagent (PolyPlus-transfection Inc, France), with 1.5µg of either wild type (WT) or mutant TRPA1 channel. Cells were used for experiments 24 to 72 hours after transfection. In PIP_2_ experiments, an equal concentration of DrVSP (Voltage-sensitive phosphatase) and TRPA1 (WT or M978V) plasmids was used to transfect the cells.

### Plasmids and site-directed mutagenesis

Human TRPA1 cDNA was cloned into a modified pcDNA3 expression vector containing downstream IRES and dsRED2 sequences (*TRPA1*-IRES-DsRED, kind gift from Dr Iulia Blesneac and Dr James Cox). The single aminoacid variant was introduced using QuikChange II XL site-directed mutagenesis kit (Agilent).

### Electrophysiology

Whole-cell patch clamp recordings were performed using an Axopatch 200B amplifier, the Digidata 1550B Low Noise Data Acquisition System, and pClamp software suite (Molecular Devices). Data were low-pass filtered at 5 kHz and sampled at 20 kHz. Capacity transients were cancelled and series resistance compensated at 70% to 90%. All recordings were performed at room temperature (22°C). Current density and voltage dependence were assessed in response to a voltage-step protocol (from −100mV to +180mV, 400ms, in the whole-cell configuration), and a second pulse to −70mV (400ms), to study tail current amplitudes and derive half maximal activation potential (V_50_) and slope factors. Only positively transfected cells were considered for experiments.

The extracellular solution contained [in mM]: NaCl 127, KCl 3, CaCl_2_ 2.5, MgCl_2_ 1, HEPES 10 and Glucose 10. pH was adjusted to 7.3 with NaOH.

For nomically Calcium-free conditions, the extracellular solution contained [in mM]: NaCl 140, KCl 3, EGTA 1, MgCl_2_ 1.3, HEPES 10 and Glucose 20. pH was adjusted to 7.3 with NaOH. Patch pipettes had a typical resistance of 3-5MΩ. Patch pipettes were pulled from borosilicate glass capillaries (1.5 mm outer diameter, 0.84 mm inner diameter; World Precision Instruments). The intracellular solution contained [in mM]: KCl 130, CsCl 5, EGTA 10 and HEPES 10. A holding potential of −60 mV was applied for all protocols. Voltage-ramp protocol was used (from −100mV to +100mV, 500ms duration, applied ever 5 sec) to assess I-V curves in absence of extracellular calcium. Current voltage curves (I-V curves) were fitted using a combined Boltzmann and linear ohmic relationship, described as: I/I_max_ = G_max_ (V_m_ – E_rev_) / (1+ exp^(V50-Vm/k)^). The reversal potential (E_rev_) was extrapolated from the fitting.

The normalised conductance-voltage (G-V) curves for activation were fitted with a Boltzmann eq. described as: G/G_max_ = {1+exp[V-V_50_)/k]}^-1^. Tail currents were fitted with non-linear regression within GraphPad Prism using the Boltzmann equation: I/I_max_ = 1 / (1+ exp[-(V-V_50_)/k]). V_50_ represents the membrane potential at half-maximal channel (in)activation in mV, V_m_ is the membrane voltage in mV and k is the slope factor. The goodness of fit was R2 > 0.9 for all Boltzmann fits. V_50_ and slope (k) values were extrapolated from Boltzmann fits of tail currents measured immediately after stepping back to a fixed potential (−70mV), when the number of open channels equals the steady-state open probability at the preceding test voltage. In this case, the tail amplitude is proportional to open probability (Po) and therefore to conductance (G) at that voltage, and the driving force (V – E_rev_) is constant.

### Immunocytochemistry

HEK293T cells were transiently transfected with either hTRPA1-IRES-DsRed or hM978V TRPA1-IRES-DsRed as described above. Cells were exposed to either AITC (25µM) or Menthol (50µM) for different incubation times, either in calcium-free or calcium-containing ECS. The calcium-containing ECS contained in [mM]: NaCl 127, KCl 3, CaCl_2_ 2.5, MgCl_2_ 1, HEPES 10, Glucose 10; pH was adjusted to 7.3 with NaOH. The calcium-free ECS contained in [mM]: NaCl 140, KCl 3, EGTA 1, MgCl_2_ 1.3, HEPES 10, Glucose 20; pH was adjusted to 7.3 with NaOH.

After treatment, cells were fixed with 4% paraformaldehyde (PFA) in PBS for 10–15 minutes at room temperature. Following two washes with PBS, cells were incubated with wheat germ agglutinin (WGA–Alexa Fluor 555 conjugate; 1 µg/mL, ThermoFisher Scientific) to label the plasma membrane, according to the manufacturer’s instructions. After two additional PBS washes, cells were permeabilized and blocked for 30 minutes at room temperature in PBS containing 0.1% Triton X-100 and 5% (v/v) normal goat serum. To detect surface expression of TRPA1 (wild-type or mutant channels), cells were incubated overnight at 4 °C with a mouse monoclonal anti-TRPA1 antibody (ANKTM1, clone C5; 1 µg/mL, sc-376495, Santa Cruz Biotechnology). The following day, cells were washed three times with PBS, under gentle agitation, and incubated with a goat anti-mouse Alexa Fluor 488 secondary antibody (1:500, ThermoFisher Scientific). After three final PBS washes, under gentle agitation, coverslips were mounted using Vectashield antifade mounting medium (VectorLabs).

### Image Acquisition and Data Analysis

Images were acquired using a Zeiss LSM-700 confocal microscope and processed with a ZEN software. Image analysis was performed using FiJi (ImageJ plugin, NIH).

In brief, as previously described [42, 68], the mean surface fluorescence intensity of each cell was quantified by manually selecting a region of interest (ROI) encircling the cell surface. Wheat germ agglutinin (WGA) membrane staining was used to delineate the plasma membrane and define a juxtamembrane ROI, which was then superimposed on the TRPA1 fluorescence image to measure surface channel expression. A separate intracellular ROI, excluding the nucleus, was used to determine the mean cytoplasmic background fluorescence intensity. The background fluorescence was subtracted from the mean surface fluorescence intensity, and the ratio of membrane-to-cytoplasmic TRPA1 expression was calculated for each cell and averaged across conditions.

Imaging parameters, including laser power, gain, and objective magnification, were kept consistent across conditions to enable direct comparison between experimental groups. Images for all experimental groups were analysed using identical parameters and image analysis was performed in a blinded manner. For each condition, data were averaged across at least two independent transfections, with a minimum of 5 positively transfected cells analysed per transfection.

### Statistical Analysis

Data are presented as mean ± standard error of the mean (S.E.M.), unless otherwise specified. Independent two-tailed t-test was used for comparisons between two groups, and two-way ANOVA and a Tukey post-hoc test for multiple-group comparison. Normality was assessed using the Shapiro–Wilk test, and non-parametric tests were applied when normality criteria were not met. Statistical analyses were performed using Excel, Origin, or GraphPad Prism (GraphPad Software, Inc., La Jolla, USA). All grouped data are mean ± SEM; A p value ≤ 0.05 was considered statistically significant. * p < 0.05, ** p < 0.01, *** p < 0.001; **** p < 0.0001.

### Molecular dynamic simulations

#### Coarse-grained molecular dynamics simulation

TRPA1 channel (residue 445 to 751 and residue 762 to 1051) was converted to a MARTINI2.2 coarse-grained representation using martinize.py, embedded in 90% palmitoyl-oleoyl-phosphatidylcholine (POPC) bilayer and 10% palmitoyl-oleoyl-phosphatidyl-4,5-inositolbisphosphate (PIP_2_) and solvated in CG-water and 0.15 M NaCl using insane.py. All simulations were carried out using the Martini2.2 biomolecular forcefield. The protein’s tertiary and quaternary structures were maintained by applying an elastic network with a force constant of 1000 kJ mol−1 nm−2 between two coarse-grained backbone particles within 0.5–0.9 nm. A temperature of 323 K was maintained with V-rescale temperature coupling, while 1 bar pressure was controlled using semi-isotropic Parrinello–Rahman pressure coupling. Systems were energy minimised using the steepest descents algorithm, equilibrated for 500 ns and simulations were conducted for 10 us for 5 repeats. All simulations were conducted using GROMACS-2021.5. The PIP_2_ binding site was identified using PyLIPID, VolMap and contact analysis, and then the pose nearest to M978 was selected for all-atoms simulation.

#### All-atoms molecular dynamics simulation

The coarse-grained simulation system with PIP_2_ in each binding site was converted to atomistic using the CG2AT2 pipeline. The M978V mutant was generated using mutagenesis wizard on PyMOL to avoid steric clashes. All simulations were carried out using CHARMM36m biomolecular forcefield with 2 fs timestep. The systems were energy minimised using the steepest descents algorithm, with non-hydrogen atoms restrained at 1000 kJ mol^−1^ nm^−2^. This was followed by a 5 ns equilibration for the system where the C_α_ backbone on the protein and the non-hydrogen atoms on the PIP_2_ molecules were restrained with 1000 kJ mol^−1^ nm^−2^. A temperature of 310 K was maintained with V-rescale temperature coupling, while 1 atm pressure was controlled using semi-isotropic Parrinello–Rahman pressure coupling. The production run was conducted for 500 ns for 3 repeats. All simulations were conducted using GROMACS-2021.5.

## Author Contributions

The project was conceived by M.C. and D.L.H.B. M.C. and D.H.L.B. wrote the manuscript, and all authors reviewed and edited the final version. M.C. planned, designed and executed electrophysiology and immunocytochemistry experiments. A.C.T. and M.L. assessed the human participants. T.P. designed, performed and analysed MD simulation experiments. M.C. designed, executed and analysed immunocytochemistry experiments and S.C. executed and analysed immunocytochemistry experiments. M.C. designed, planned, executed and analysed the calcium imaging experiments, and S.V.K executed and analysed the calcium imaging experiments (data not included).

## Competing Interest Statement

This research was funded in whole, or in part, by the Wellcome Trust (109915/Z/15/Z, 083259, 202747/Z/16/Z). D.L.H.B. has acted as a consultant on behalf of Oxford Innovation (Amgen, Bristows, LatigoBio, GSK, Ionis, Lilly, Olipass, Orion, Regeneron and Theranexus). He has received research funding from Lilly, and an industrial partnership grant from the BBSRC and AstraZeneca. A.C.T. is supported by MRC Clinician Scientist Fellowship (MR/Z504075/1). A.C.T. and D.L.H.B. are supported by the MRC/Versus Arthritis funded PAINSTORM consortium, which is part of the Advanced Pain Discovery Platform (MR/W002388/1). A.C.T. has acted as a consultant on behalf of Oxford Innovation (Vertex Pharmaceuticals).

## Acknowledgments

We thank all the subjects who took part in the study. The study was approved by the East of England Cambridge South National Research Ethics Committee (REC), reference 13/EE/0325. We thank Dr Iulia Bleasneac for her valuable comments and insights on the experimental design of the electrophysiological recordings. We thank Dr Gustavo Chavez Barboza for helpful discussion and insight regarding the PIP_2_ experiments. We thank NTU-HPC for the computational support.

---

**Supplementary Figure 1.**
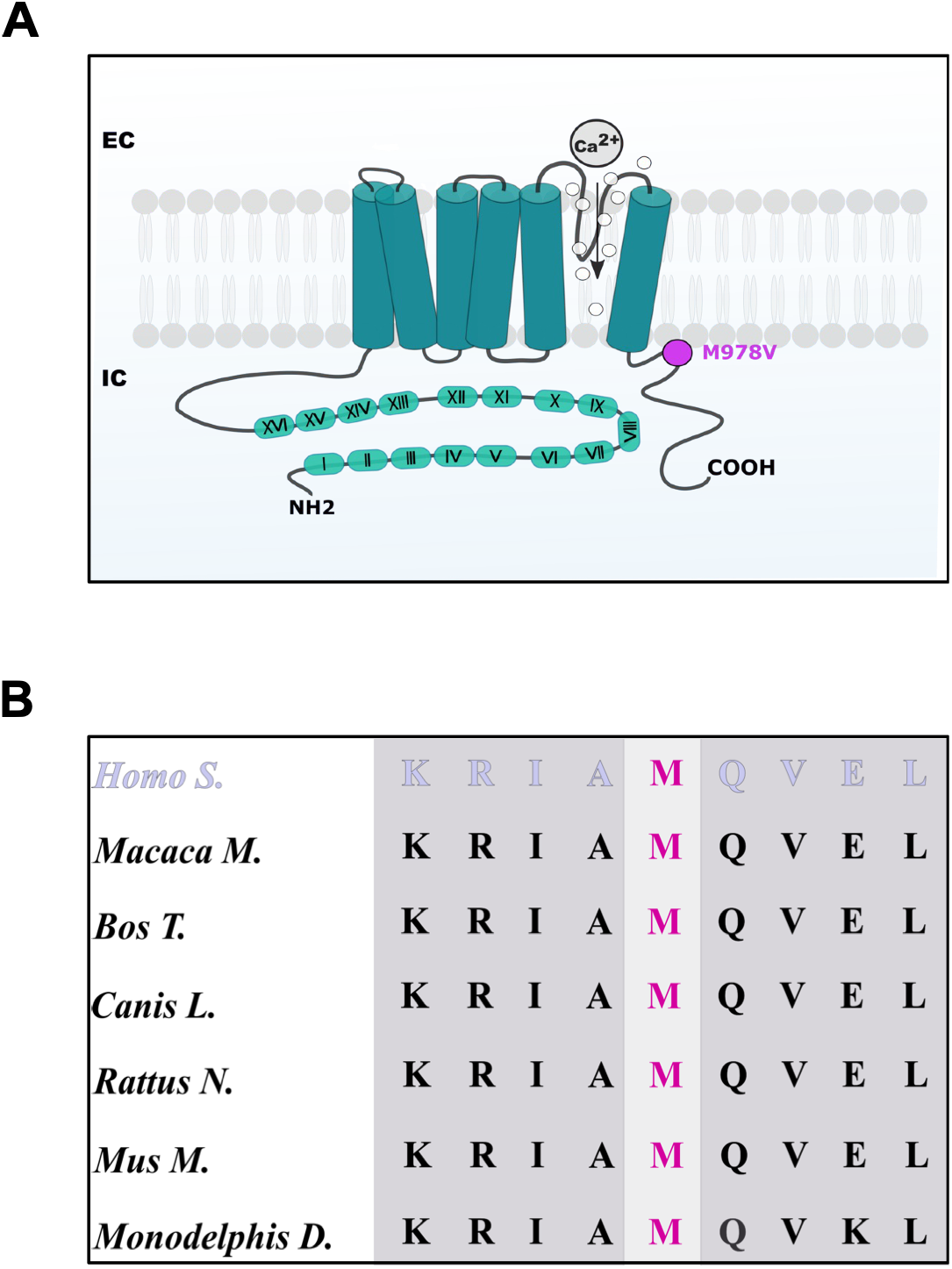
The amino-acid M978 is highly conserved across mammalian species. Legend: **(A)** In the diagram of the human TRPA1 channel, the novel genetic variant M978V (a Valine substituting Methionine in position 978) is located in the Carboxyl-terminal region of the channel and is depicted in magenta. (**B**) The mutated amino-acid Methionine is highly conserved across different mammalian species.

**Supplementary Figure 2.**
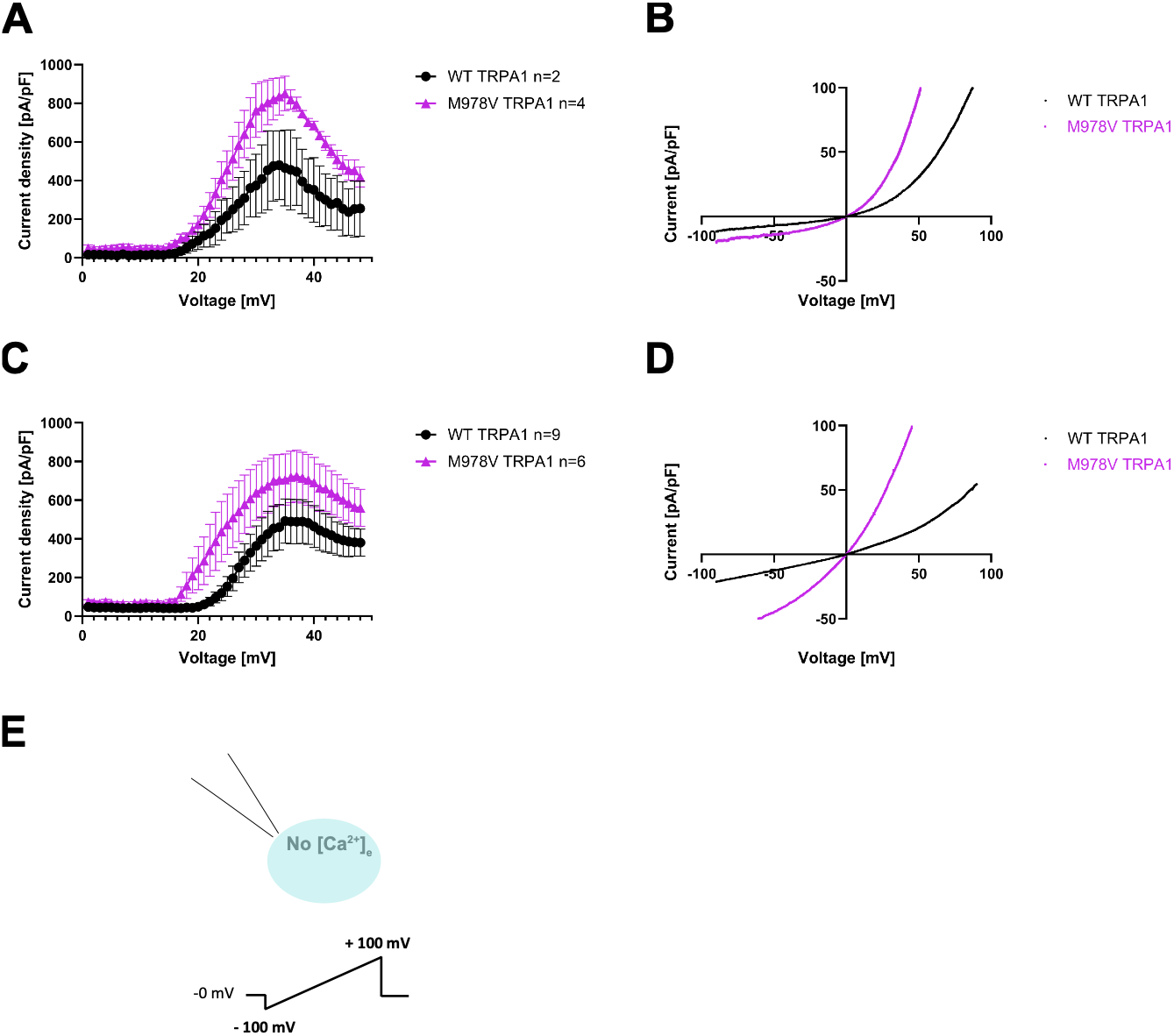
hTRPA1 M978V produces a concentration-dependent linearisation of the I-V curve in response to AITC and in absence of Ca^2+^_Ext_. Legend: Panels **A** and **C** show current density (extrapolated at positive potentials) for either WT or mutant channels (in black and magenta, respectively). Representative I-V curves are shown in panel **B** and **D**, at different AITC concentrations (25 and 100µM, respectively). The I-V relationship shows increased linearisation at higher doses of AITC. The applied voltage-ramp protocol and the experimental conditions (absence of both intracellular and extracellular calcium) are depicted in the diagram in **E**.

**Supplementary Figure 3.**
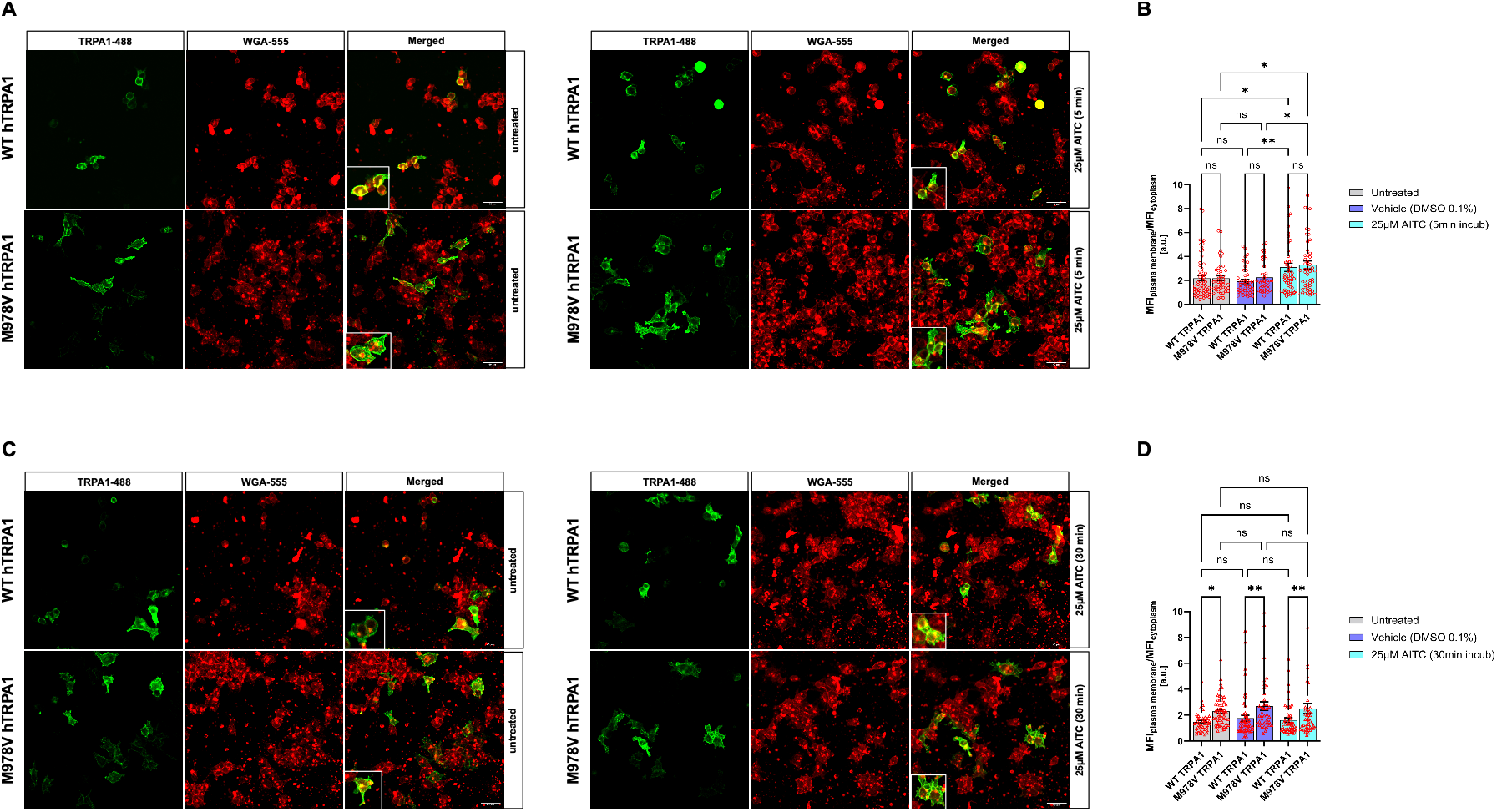
Surface membrane expression of TRPA1 M978V is enhanced after long exposure to AITC in the presence of Ca^2+^_Ext_. Legend: Representative images of HEK293T cells stained with wheat germ agglutinin (WGA) to label the plasma membrane and immunostained with an anti-TRPA1 antibody are depicted in **(A)** and **(C)**. **(B)** and **(D)** reports quantification of cell surface expression of WT and mutant TRPA1 channels following treatment with either ECS, vehicle or 25µm AITC after 5 minute incubation (untreated WT n = 65, M978V n = 44, DMSO WT n = 44, M978V n = 40, AITC WT n = 59, M978V n = 53) or 30 min incubation (untreated WT n = 40, M978V n = 55, DMSO WT n = 53, M978V n = 39, AITC WT n = 51, M978V n = 46), respectively, in the presence of extracellular calcium. Scale bar: 40µm. All grouped data are mean ± SEM, * p < 0.05, ** p < 0.01, *** p < 0.001; **** p < 0.0001.

**Supplementary Figure 4.**
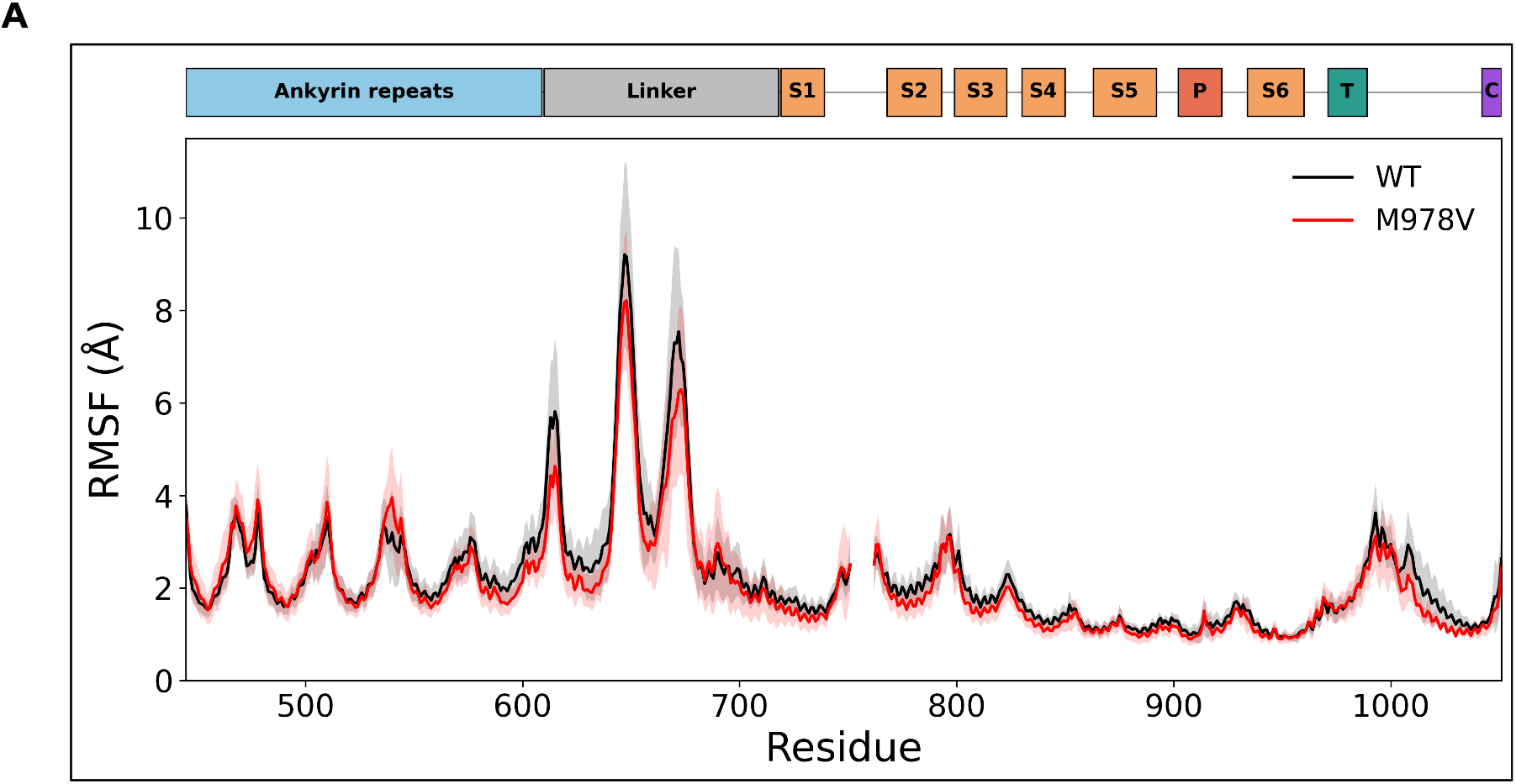
Protein flexibility and stability are not altered by the M978V mutation. Legend: (**A**) C-alpha atom root-mean-square fluctuation (RMSF) of human TRPA1, comparing wild-type (WT, black) and the M978V variant (red). RMSF from all-atoms MD simulations were averaged over 12 independent measurements per residue (4 subunits of the homotetramer × 3 replicate trajectories); the shaded region shows ±1 standard deviation. The bar above the plot maps the domain topology (UniProt O75762): ankyrin repeats, linker, transmembrane helices S1–S6, pore region (P), TRP helix (T), and C-terminal coiled coil (C) region.

